# Optogenetic stimulation of dorsal striatum bidirectionally controls seizures

**DOI:** 10.1101/2024.09.18.613710

**Authors:** Safwan K. Hyder, Willian Lazarini-Lopes, Jonathan Toib, Gabrielle Williams, Alex Sukharev, Patrick A. Forcelli

## Abstract

Engagement of the striatum (caudate/putamen) and other basal ganglia nuclei during seizures was first observed over 75 years ago. Basal ganglia output nuclei, and the substantia nigra pars reticulata, in particular, have well-established anti-seizure effects across a large array of experimental models. However, striatal control of seizures is understudied. To address this gap, we used optogenetic approaches to activate and inactivate neurons in the dorsal striatum of Sprague-Dawley rats submitted to the gamma-butyrolactone (GBL) model of absence epilepsy, amygdala kindling model of temporal lobe epilepsy, and pilocarpine-induced Status Epilepticus (SE). All tests were performed on a within-subject basis. Animals were tested in two different light frequencies (5 Hz and 100 Hz). Open-loop (continuous light delivery) optogenetic activation of the dorsal striatal neurons robustly suppressed seizures in all models. On the other hand, optogenetic silencing of the dorsal striatal neurons increased absence seizure expression and facilitated SE onset but had no effect on kindled limbic seizures. In the GBL model, we also verified if the closed-loop strategy (light delivery in response to seizure detection) would be enough to induce antiseizure effects. On-demand light delivery in ChR2-expressing animals reduced SWD duration, while the same approach in ArchT-expressing animals increased SWD duration. These results demonstrated previously unrecognized anti-absence effects associated with striatal continuous and on-demand neuromodulation. Together, these findings document a robust, bidirectional role of the dorsal striatum in the control of seizure generation and propagation in a variety of seizure models, including focal seizure onset and generalized seizures.

## Introduction

Almost 75 years ago, Stoll and colleagues demonstrated engagement of striatum (caudate/putamen) during seizures triggered by stimulation of the temporal lobe in cats (1). This work, followed by that of Udvarhayli and Walker, demonstrated engagement of multiple basal ganglia nuclei, including the striatum and the substantia nigra, in focal onset seizures in monkeys (2). However, the basal ganglia remain most commonly associated with their role in movement control and reward and reinforcement. Accordingly, the role of these structures in the initiation, propagation, and generalization of seizures has been a lingering question in the field. Significant work in the 1980s and 1990s demonstrated that focal inhibition of the basal ganglia output nucleus, the substantia nigra pars reticulata (SNpr), potently suppresses seizures in a wide range of experimental models (3–6). More recently, we demonstrated that this effect is mediated by projections from the SNpr to the deep and intermediate layers of the superior colliculus (DLSC) (7–9). However, the upstream regulators of the SNpr remain poorly studied in the context of epilepsy. In a series of studies, Turski and colleagues found that glutamate agonist or GABA antagonist injection into the striatum suppresses seizures in both the amygdala kindling model and the pilocarpine model of temporal lobe epilepsy (10, 11). More recently, Brodovskaya and colleagues (12) found, using a genetic activity mapping strategy, that a subpopulation of neurons within the striatum, those that project to the globus pallidus external segment and express dopamine type 2 (D2) receptors were preferentially activated by focal onset motor seizures as compared to dopamine type 1 (D1) receptors expressing neurons which project to and inhibit the SNpr. However, the impact of reversible inhibition of the striatum on seizures is unknown and, unlike downstream components of the extended basal ganglia (i.e., the SNpr, the DLSC), the role of striatal stimulation in gating other forms of seizures (e.g., absence seizures) still needs to be properly investigated.

While focal pharmacology is a powerful tool for probing brain circuit function, it is limited by potential drug diffusion and does not allow for real-time control of neuronal activity. To address this gap, we employed bi-directional optogenetic manipulations of the dorsal striatum to either activate or silence the striatum neurons. We tested our striatal manipulations in three preclinical seizure models. The first was the gamma-butyrolactone (GBL) evoked spike-and-wave discharges (SWDs) which models absence epilepsy. Here we systemically administered GBL while recording cortical EEG and subsequently modulated the striatum using both open-loop (continuous) and closed-loop (responsive, or on-demand neurostimulation) stimulation. The second model was the amygdala kindling model of temporal lobe epilepsy. Here we performed our dorsal striatal manipulations with subsequent delivery of electrical stimulus to evoke temporal lobe-like seizures. For our third model, we utilized the pilocarpine model of status epilepticus (SE), in which we performed dorsal striatal manipulations to test their effects on the latency to pilocarpine-induced SE.

We found that in the GBL model of absence seizures, striatal manipulations elicited bidirectional control of seizures. Striatal activation resulted in a reduction of SWD activity, while striatal silencing exacerbated SWDs. Furthermore, using a closed-loop approach in which optogenetic manipulation of the striatum was only delivered in response to detected SWDs, we found a reduction and an increase in SWD duration in response to striatal activation and silencing, respectively. In the amygdala kindling model, we found a unidirectional effect in which striatal activation resulted in a reduction of seizure severity and duration. Finally, in the pilocarpine model of SE, we again found a bidirectional effect: striatal activation increased the latency to SE onset, while the opposite effect was induced with striatal silencing. Together, these data demonstrate that the dorsal striatum provides strong bidirectional control over seizure activity and demonstrate that striatal neurostimulation can be deployed to control seizures in both open- and closed-loop neurostimulation paradigms.

## Results

### Histological validation

Fig. 1 shows the histological confirmation of opsin expression in dorsal striatum neurons. Fig. 1A shows a representative case injected with AAV5-hSyn-ChR2(H134R)-mCherry for activation experiments and Fig. 1B shows cell bodies of ChR2-expressing neurons. Fig. 1C shows a representative case injected with AAV8-CAG-ArchT-GFP for inhibition experiments and Fig. 1D shows cell bodies of ArchT-expressing neurons. Fig. 1E and 1F show mapping of fiber optic tips across cases in the GBL and kindling experiments, respectively. Note in both the activation and inhibition experiments, some labeling along the deep cortical layers was noted, which may have been a result of virus reflux up the needle track and/or retrograde transport of AAV (13). In either case, it is unlikely to have contributed to the effects we report below, as the fiber optics were placed in the dorsal striatum, ventral to the off-target expression. Of the 92 animals used in this study, 11 were excluded for insufficient virus expression or misplaced fiber optics. Histological determination was made while blinded to seizure data. All cases were reviewed by either SH and PAF or WLL and PAF.

**Figure 1.**
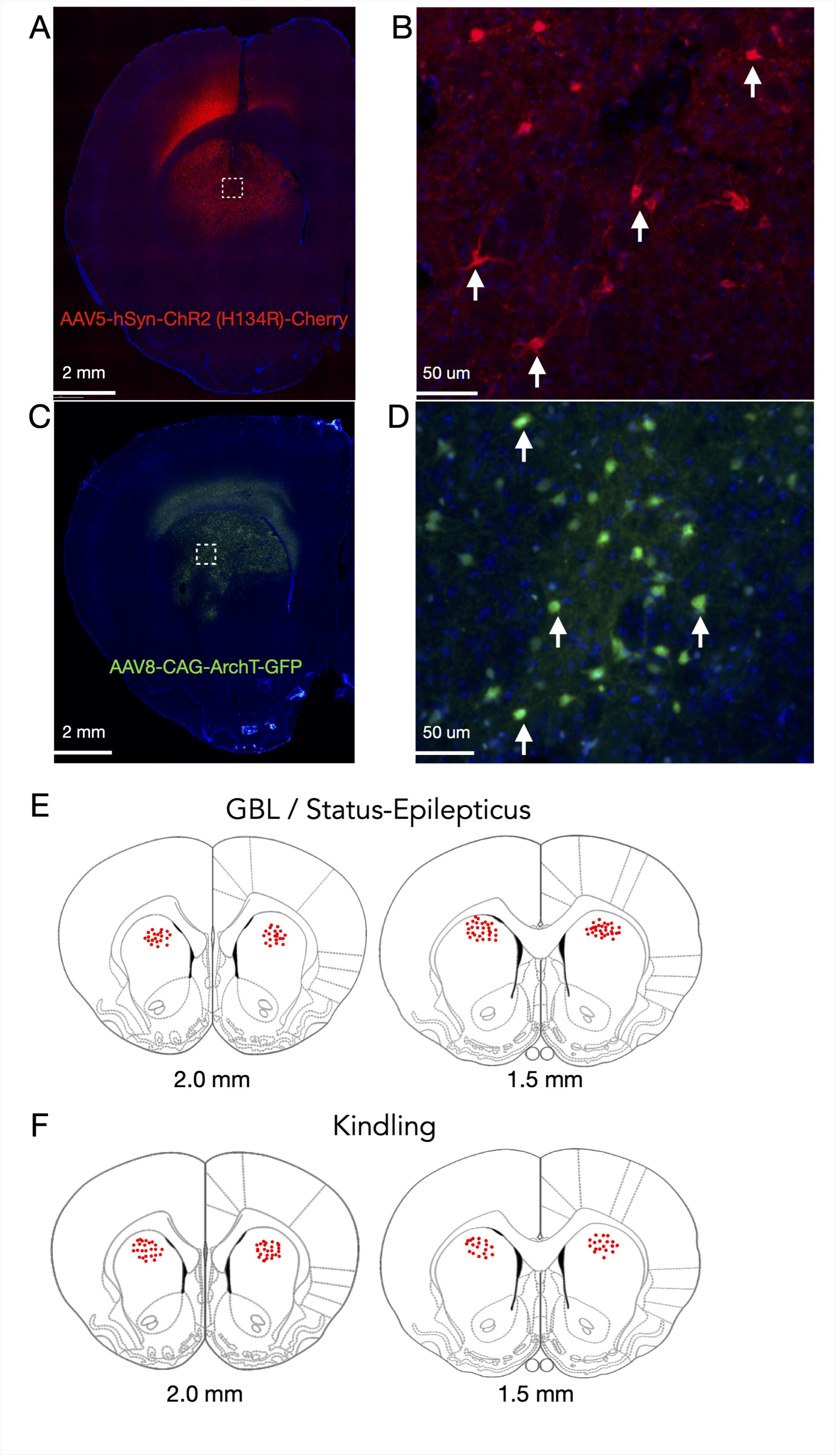
Histological verification of virus expression into the dorsal striatum. (A) Representative image of ChR2-mcherry expression into the striatum (1.6x magnification, mcherry fluorescence in red). (B) Cell bodies of neurons transfected with AAV5-Syn-ChR2 (H134R)-mcherry expressing ChR2 (20x magnification). Arrows indicate ChR2-mcherry positive cells. (C) Representative image of ArchT-GFP expression into the striatum (1.6x magnification, GFP fluorescence in green). (D) Cell bodies of neurons transfected with AAV8-CAG-ArchT-GFP expressing ArchT-GFP (20x magnification). Arrows indicate ArchT-GFP positive cells. (E-F) Atlas planes modified from Paxinos and Watson (44) with coronal sections of the striatum. Red dots represent the place of fiber optics’ tips into the dorsal striatum from 1.5 mm to 2.0 mm anterior to bregma in animals submitted to the GBL protocol of absence seizures and to the kindling protocol of temporal lobe epilepsy.

### Optogenetic open-loop activation of dorsal striatal neurons suppresses absence seizures

Fig. 2 shows the results from open loop optogenetic activation of the striatum in ChR2- expressing animals. Animals were injected with GBL (100 mg/kg) and cortical EEG activity was monitored for 30 min. Animals were tested on a within-subject basis with one session without light delivery and one session with light delivery (either 5 Hz or 100 Hz); this sequence was repeated for the second frequency of light delivery.

**Figure 2.**
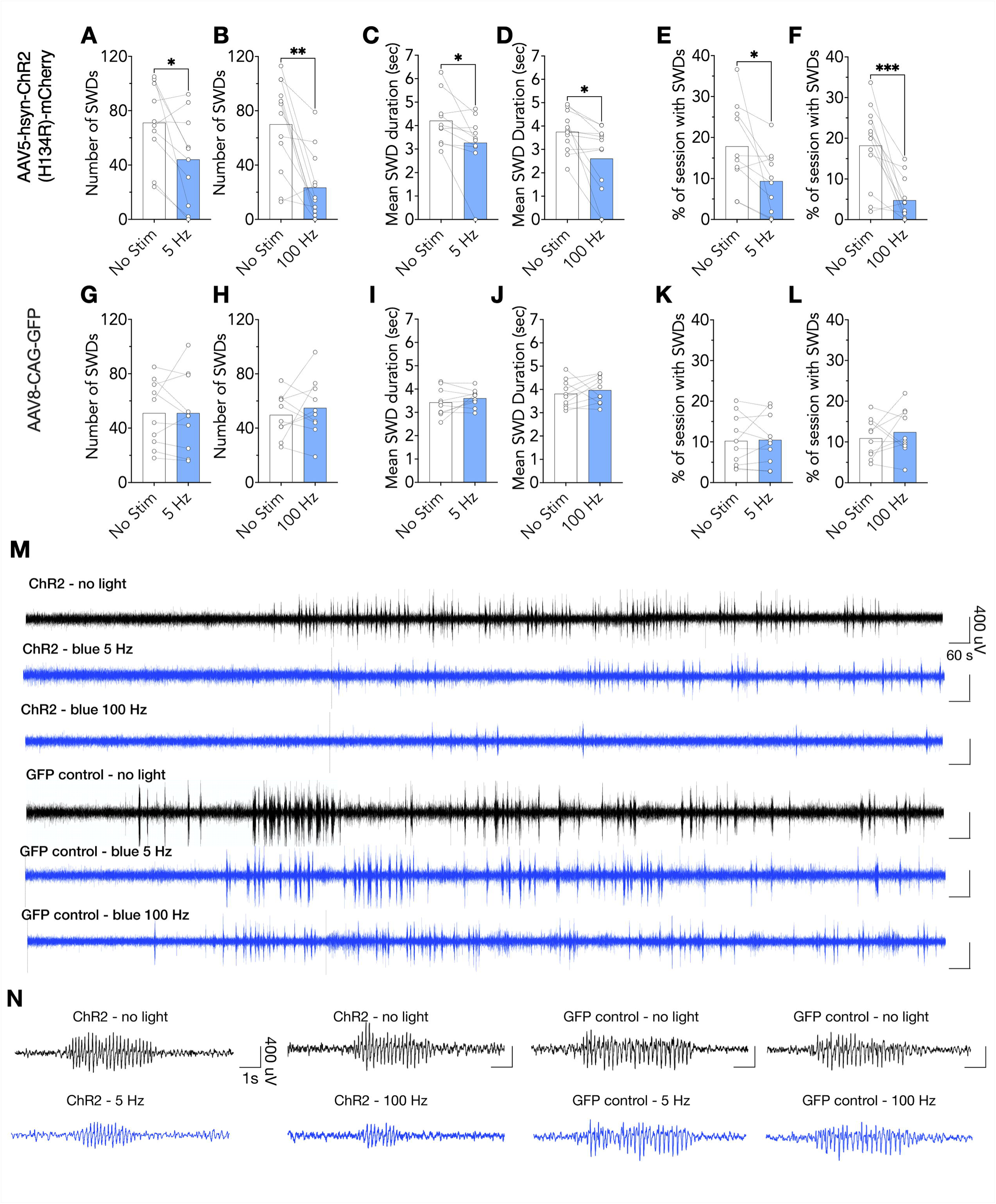
Open loop optogenetic activation of dorsal striatal neurons suppresses absence seizures. (A-F) Number of spike-and-wave discharges (SWDs), SWD mean duration, and percentage of time seized in ChR2 animals submitted to blue light open loop optogenetic stimulation under 5 Hz and 100 Hz. Optogenetic activation of striatal neurons under blue lights 5 Hz (n=10) and 100 Hz (n=12) frequencies attenuated SWD manifestation in ChR2 animals, reducing the number, duration, and percentage of time displaying SWDs. (G-L) Number of SWDs, SWD mean duration, and percentage of time seized in GFP control animals (no opsin) submitted to blue light stimulation with 5 Hz (n=10) and 100 Hz (n=10). None of the frequencies of stimulations modified SWDs in GFP control animals. (M) Representative EEG traces from ChR2 and GFP animals submitted to 30 min cortical EEG recordings under no light (black traces), 5 Hz and 100 Hz stimulations (blue traces). (N) Representative SWDs from ChR2 and GFP control animals submitted to non-stimulated test session (black traces) and to optogenetic activation of the striatum under 5 and 100 Hz (blue traces). Significance: p<0.05. *p<0.05, **p<0.01, ***p<0.001.

5 Hz optogenetic activation of the striatum significantly decreased the number of SWDs during the test session as compared to the no stimulation session (Fig. 2A, paired t-test, t=2.656, df=9, p=0.0262). We observed a similar response in animals tested with 100 Hz light delivery (Fig. 2B, paired t-test, t=4.326, df=11, p=0.0012). In addition to the reduction in the number of SWDs, optogenetic activation of the striatum also reduced the mean SWD duration with 5 Hz (Fig. 2C, paired t-test, t=2.656, df=9, p=0.0262) and 100 Hz light delivery (Fig. 2D, paired t-test, t=3.020, df=11, p=0.0116). Consistent with both of these findings, the percentage of the session with SWD activity (a composite measure), was reduced by both 5 Hz (Fig. 2E, paired t-test, t=2.550, df=9, p=0.0312) and 100 Hz (Fig. 2F paired t-test, t=4.587, df=11, p=0.0008) stimulation.

In opsin-negative control animals (expressing just green fluorescent protein - GFP), blue light delivery at either 5 or 100 Hz frequencies had no effect on SWD frequency (Fig. 2G and H, p>0.05), duration (Fig. 2I and J, p>0.05), and percentage of time seized during the session (Fig. 2K and L, p>0.05). Figure 2M shows representative EEG traces from animals submitted to open-loop optogenetic activation of dorsal striatum (30 min duration). Fig. 2N shows representative SWDs displayed by ChR2 and GFP animals (5 and 100 Hz) and non-stimulated test sessions. Assessment of cortical EEG power spectra normalized to peak power showed no effects of optogenetic manipulation on cortical EEG spectra activity in both ChR2 and GFP animals tested with blue light under no seizure conditions (*SI Appendix*, Fig. S1-S3).

These data demonstrate a previously unknown anti-absence effect of striatal stimulation and suggest that striatal stimulation may provide the same broad-spectrum anti-seizure effect previously described for the SNpr and DLSC (7–9).

### Optogenetic open-loop inhibition of dorsal striatal neurons exacerbates absence seizures

Given that activation of the striatum suppressed SWDs, we next sought to determine if this effect was bidirectional. Thus, we optogenetically inhibited the dorsal striatum and again assessed SWD burden in the GBL model. Optogenetic inhibition of the striatum significantly increased the number of SWDs (Fig. 3A, paired t-test, t=3.461, df=8, p=0.0086). By contrast, we found no difference in mean SWD duration (Fig. 3B, paired t-test, t=1.473, df=8, p=0.1789), despite an increase in SWD duration in all but one animal. The percentage of time seized (Fig. 3C, paired t-test, t=4.435, df=8, p=0.0022) was significantly increased, likely driven by the increased number of SWDs.

**Figure 3.**
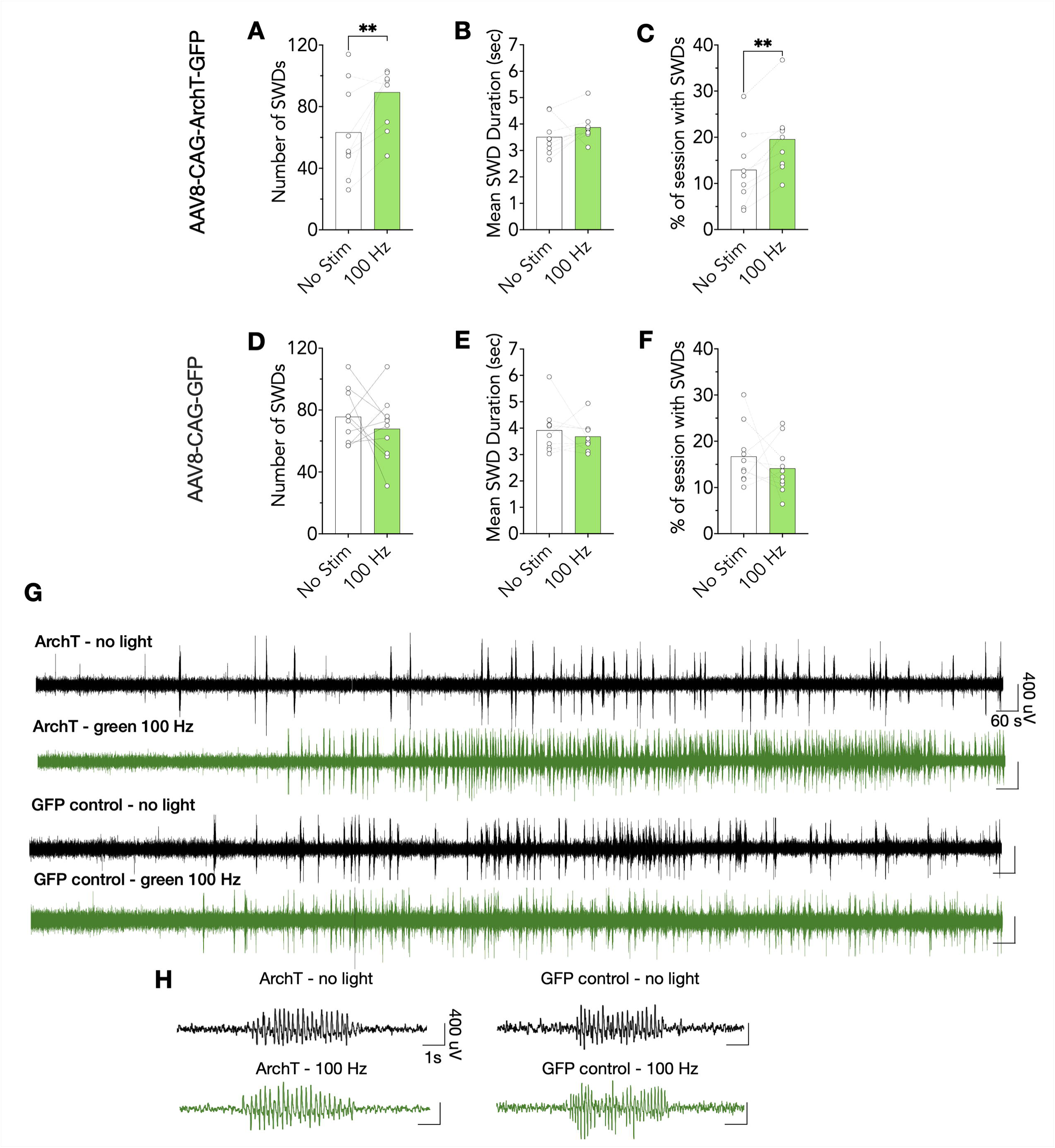
Open loop optogenetic inactivation of dorsal striatal neurons increases absence seizures. (A-C) Number of spike-and-wave discharges (SWDs), SWD mean duration, and percentage of time seized in ArchT animals (n=9) submitted to test sessions with no stimulation and with green light open loop optogenetic silencing under 100 Hz. Optogenetic silencing of striatal neurons increased SWD manifestation in ArchT animals, with a significant increase in the number of SWDs and the percentage of time seized, but with no significant alteration for SWD duration. (D-F) Number of SWDs, SWD mean duration, and percentage of time seized in GFP (no opsin) control animals (n=10). No alteration was detected for SWD manifestation in GFP animals tested under green light 100 Hz and no light. (G) Representative EEG traces from ArchT and GFP animals submitted to 30 min of cortical EEG recordings under no light (black traces) and green light 100 Hz (green traces). (H) Representative SWDs from ArchT and GFP control animals submitted to non-stimulated test session (black traces) and to optogenetic silencing of the striatum under 100 Hz (green traces). Significance: p<0.05. *p<0.05, **p<0.01.

In opsin-negative control animals, green light delivery was without effect on SWD frequency (Fig. 3D, p<0.05), SWD mean duration (Fig. 3E, p<0.05), and percentage of time seized (Fig. 3F, p>0.05). Fig. 3G shows representative EEG traces from rats during open-loop optogenetic inactivation of dorsal striatal neurons and Fig. 3H shows representative SWDs from ArchT and GFP animals. Similarly to ChR2 animals, analysis of EEG power spectra showed no effect of optogenetic inactivation of the striatum on cortical EEG spectra in either ArchT and GFP animals tested under no seizure conditions (*SI Appendix*, Fig. S1-S3).

Together with our results using optogenetic activation (Fig. 2), these results demonstrate that striatal activity bidirectionally modulates absence-like seizures in the GBL model. These data provide proof-of-concept for optogenetic neuromodulation of the striatum to control seizures. However, given the importance of the striatum for movement control and cognitive functions, we explored the efficacy of using a more tailored neurostimulation approach that limited stimulation to periods of seizure activity. Thus, we next sought to determine if closed-loop (on-demand) neuromodulation would suppress seizures, as has been reported in hippocampus, medial septum, and cerebellum in models of temporal lobe epilepsy (14–18), as well as in superior colliculus in models of absence seizures (8, 9).

### Optogenetic closed-loop neuromodulation of dorsal striatal neurons controls absence seizures

We performed online seizure detection and randomized half of detected events to receive light delivery (5 second light duration) and half to serve as control events. As shown in Fig. 4, closed-loop stimulation at 5 Hz did not terminate SWDs (Fig. 4A, paired t-test, t=0.3609, df=7, p=0.7288), but closed-loop stimulation at 100 Hz did (Fig. 4B, paired t-test, t=5.453, d=7, p=0.0010). Interestingly, given that open-loop inhibition for the striatum did not significantly impact SWD duration, closed-loop inhibition at 100 Hz significantly increased SWD duration (Fig. 4C, paired t-test, t=8.796, df=5, p=0.0003). Consistent with these findings, we found a left-shift in the cumulative frequency distribution for seizure duration with 100 Hz optogenetic stimulation (Fig. 4E, Kolmogorov-Smirnov test, p<0.0001), but no difference with 5 Hz (Fig. 4D, p>0.05). In contrast to the decreased seizure duration observed with optogenetic stimulation in ChR2-expressing animals, we observed a right-shift in the cumulative frequency distribution for seizure duration with 100 Hz inactivation in ArchT-expressing animals (Fig. 4F, Kolmogorov-Smirnov test, p=0.0281).

**Figure 4.**
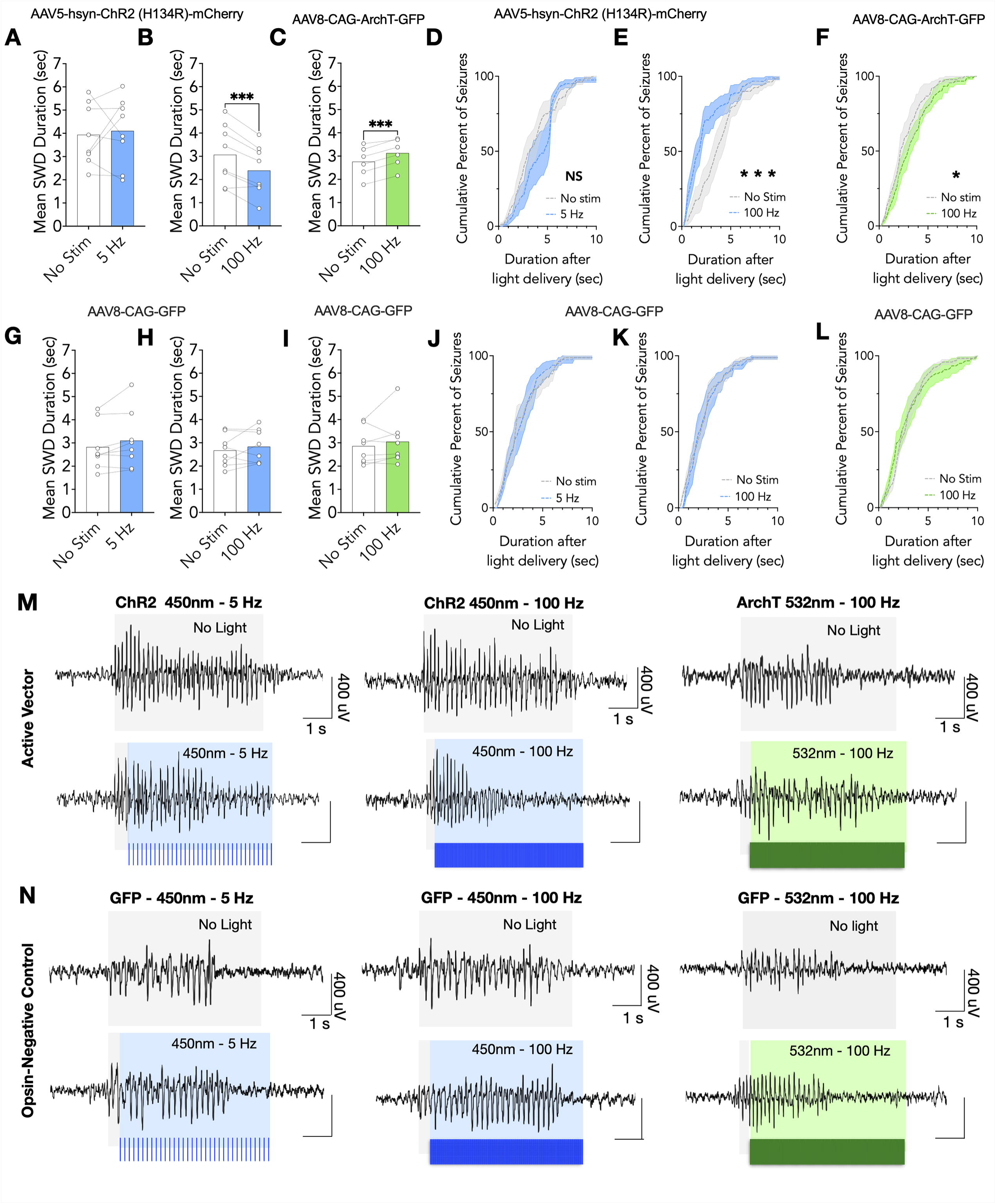
Closed-loop optogenetic activation and inactivation of dorsal striatal neurons has bidirectional effects on SWD duration. (A-B) Optogenetic closed loop activation of the dorsal striatum (blue light 100 Hz) in ChR2 animals (n=8) resulted in faster termination of SWDs when compared to non-stimulated seizures. Closed loop optogenetic activation of dorsal striatum (blue lights 5 Hz) did not modify SWDs mean duration in ChR2 animals (n=8). (C) Optogenetic closed loop silencing of the dorsal striatum in ArchT animals (green lights 100 Hz) resulted in longer seizures in comparison to non-stimulated seizures (n=6). (D-F) Cumulative distribution of SWD duration after light delivery at each frequency. Plots show the mean and 95% confidence interval for the average cumulative frequency distribution. SWD cumulative distribution duration in ChR2 animals tested under blue lights 100 Hz were shifted to the left, indicating rapid SWDs termination after light delivery in comparison to non-stimulated seizures (n=8). The opposite effect was observed in ArchT animals tested under green light 100 Hz (n=6), with a cumulative distribution shifted to the right side in comparison to non-stimulated seizures, indicating that seizures became longer after light delivery in ArchT animals. (G-I) Optogenetic closed loop stimulation (blue light 5 and 100 Hz and green light 100 Hz) of the striatum did not change SWD duration in GFP control animals (n=8). (J-L) Light delivery did not change cumulative distribution of SWD duration in GFP (no opsin) control animals (n=8) submitted to closed loop optogenetic stimulation (blue light 5 and 100 Hz and green light 100 Hz) of dorsal striatum. Plots show the mean and 95% confidence interval for the average cumulative frequency distribution. (M-N) Representative SWDs from ChR2, ArchT and GFP control rats submitted to closed loop optogenetic neuromodulation of the dorsal striatum. Grey shading shows the onset of seizure detection followed by 5 s of post-detection window without light delivery. Blue and green shading indicates the period of light delivery (5 s duration) following seizure detection on optogenetic stimulated seizures. Significance: p<0.05. *p<0.05, **p<0.01, ***p<0.001. NS=no significance.

As expected, closed-loop optogenetic neuromodulation in GFP-expressing control animals was without effect on SWDs during blue (5 and 100 Hz) or green (100 Hz) light delivery (Fig. 4G- L, p>0.05). Fig. 4M shows representative SWDs from ChR2 and ArchT animals submitted to optogenetic closed-loop neuromodulation. Fig. 4N shows representative SWDs from GFP control animals submitted to blue (5 and 100 Hz) and green (100 Hz) light delivery.

Given that both open and closed-loop protocols suppressed SWDs in the GBL model, we next compared the efficacy of the two strategies. Since blue 5Hz was not effective in the closed-loop protocol, we only compared open- and closed-loop sessions performed with 100Hz stimulation. Mean SWD duration was reduced with closed-loop stimulation as compared to open-loop stimulation (*SI Appendix*, Fig. S4, paired t-test, t=2.485, df=7, p<0.05). Thus, while open-loop stimulation reduced the occurance of SWDs, closed-loop stimulation was more effective at early termination of SWDs.

### Optogenetic activation of dorsal striatal neurons suppresses kindled limbic seizures

Given the striking bidirectional modulation of seizure activity that we observed with optogenetic activation and inhibition of the striatum in the GBL model of absence seizures, we next sought to determine if the same pattern held true for limbic seizures evoked in the amygdala kindling model of temporal lobe epilepsy. We kindled animals as previously described (19) until they displayed stable Racine Stage 5 seizures (20), and then subjected them to optogenetic neuromodulation on a within-subject basis. Optogenetic activation of the striatum with 5 Hz light delivery failed to significantly reduce the duration of electrographic seizures (afterdischarge) (Fig. 5A; paired t-test, t=2.256, df=6, p=0.0649) or behavioral (Fig. 5D; Wilcoxon signed rank test, p=0.1250) seizures triggered by amygdala stimulation. By contrast, optogenetic activation of the striatum with 100 Hz light delivery significantly reduced both electrographic seizure duration (Fig. 5B; paired t-test, t=5.350, df=9, p=0.0005) and behavioral seizure severity (Fig. 5E; Wilcoxon signed rank test, p=0.0039). In fact, 6 of 10 animals were completely protected from electrographic seizures, and 8 of 10 animals were completely protected from behavioral seizures.

**Figure 5.**
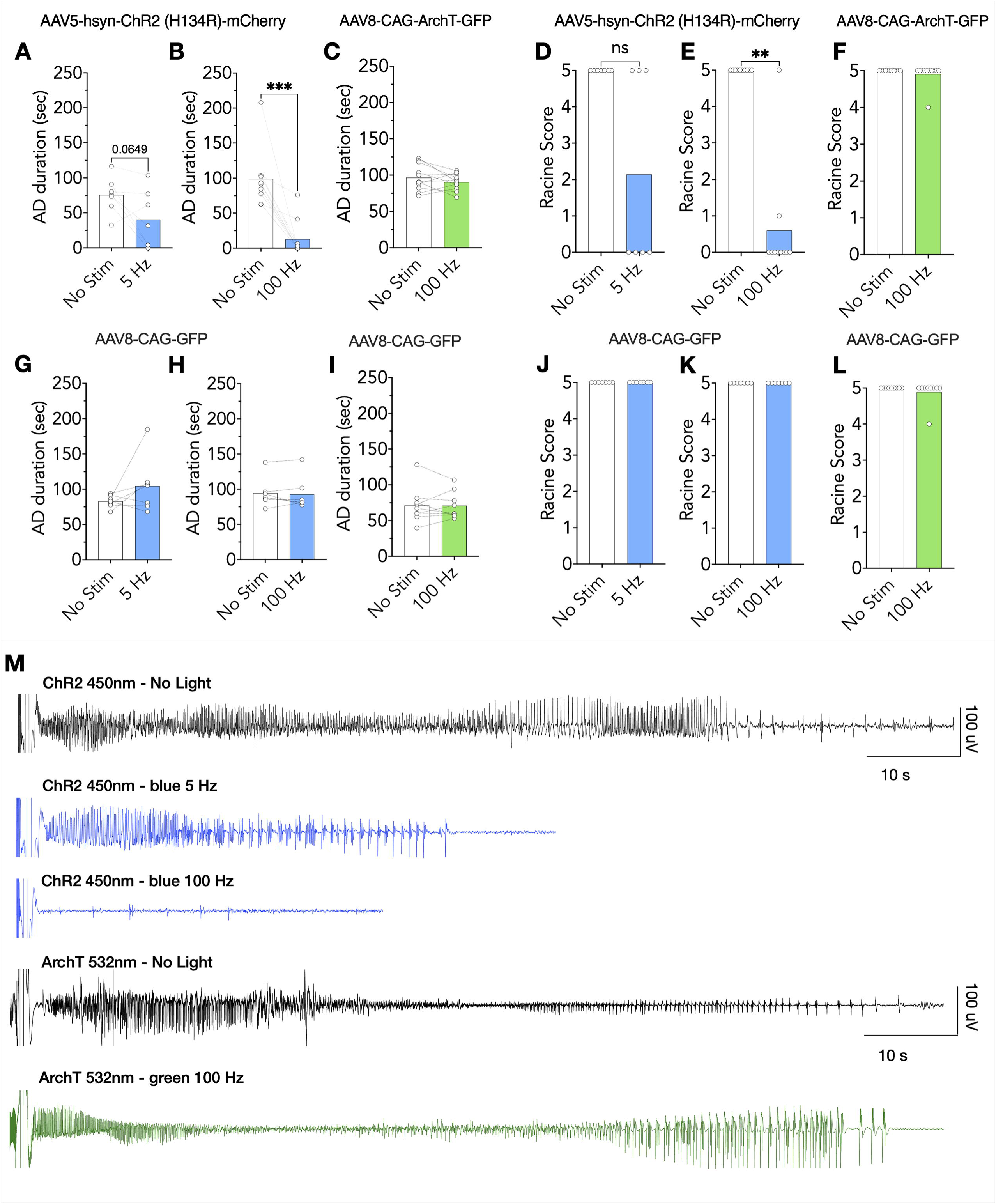
Optogenetic activation of the dorsal striatum suppressed limbic seizures in amygdala-kindled rats. (A-B) Optogenetic activation (AAV5-hSyn-ChR2 (H134R)-mCherry) of the dorsal striatum significantly reduced afterdischarge (AD) duration in ChR2 animals stimulated with blue light at 100 Hz (n=10), but only a trend towards significance was detected for ChR2 stimulated with 5 Hz (n=7). (C) Optogenetic silencing (AAV8-CAG-ArchT-GFP) of neurons from the dorsal striatum did not modify afterdischarge (AD) duration in animals stimulated with green light at 100 Hz (n=11). (D-E) Optogenetic activation of the dorsal striatum in at 100 Hz suppressed limbic seizure behavioral manifestation in fully kindled rats (n=10), but 5 Hz stimulation (n=7) had no significant effect on seizure behavior. (F) Optogenetic silencing of dorsal striatal neurons did not modify limbic seizure behavior in ArchT animals (n=11). (G-L) Optogenetic stimulation of the dorsal striatum with blue (5 and 100 Hz, n=7) and green (100 Hz, n=9) lights in GFP control animals (AAV8-CAG-GFP, opsin negative) did not modify AD duration and limbic seizure expression. Limbic seizure behaviors were scored according to the Racine scale (20). (M) Representative EEG traces from the basolateral amygdala nucleus (BLA) in ChR2 and ArchT animals during no light (black traces), blue lights (5 and 100 Hz, blue traces), and green lights (100 Hz, green traces) test sessions. Significance: p<0.05. *p<0.05, **p<0.01, ***p<0.001. NS=no significance.

Unlike the GBL model, optogenetic inactivation of the striatum was without effect on afterdischarge duration (Fig. 5C; paired t-test, t=1.307, df=10, p=0.2205), and given that a Score 5 seizure is the maximum seizure severity, a ceiling effect thus precluded a worsening of behavioral seizures (Fig. 5F; Wilcoxon signed rank test, p>0.9999). Fig. 5M shows representative EEG traces from the basolateral amygdala nucleus (BLA) activity in kindling test sessions from ChR2 and ArchT animals submitted to BLA electrical stimulation with no light and under optogenetic neuromodulation of the dorsal striatum with blue (ChR2 animals; 5 and 100 Hz) and green (ArchT animals; 100 Hz) lights.

As expected, blue (5 and 100 Hz) or green (100 Hz) light delivery were both without effect on seizure score (Fig. 5J-L; Wilcoxon signed rank test, ps>0.05) and afterdischarge duration (Fig 5G-I, ps>0.05; *SI Appendix*, Fig. S5A-B) in opsin negative kindled animals. Similar to GBL experiments, analysis of EEG power spectra of ChR2, ArchT, and GFP animals showed no effect of optogenetic neuromodulation on electrographic power in the BLA in animals tested with blue or green light under no seizure conditions (*SI Appendix*, Fig. S6).

### Optogenetic neuromodulation of dorsal striatum neurons controls pilocarpine-induced *Status Epilepticus* (SE)

Given that we did not detect a worsening of amygdala kindled seizures with optogenetic inactivation of the striatum, we sought to evaluate another model of temporal lobe seizures that displays greater sensitivity to bidirectional modulation. We turned to the pilocarpine model, which was also the model used in the early pharmacological manipulation studies of the striatum (10, 21).

For these experiments, we used a subset of animals that had completed testing in the GBL model. Animals were injected with pilocarpine (380 mg/kg) to trigger SE at least a month after the conclusion of GBL experiments. Since blue 100 Hz was the most effective frequency in both the kindling and GBL models, we only tested animals with 100 Hz light delivery. Light delivery was started 10 min before injection of pilocarpine, and we monitored the latency to SE and mortality. Animals were monitored for 90 min after pilocarpine administration.

Optogenetic activation (100 Hz) of the striatum significantly increased the latency to SE onset in the ChR2 group compared to GFP (Fig. 6A, p=0.0125 Mann-Whitney test). By contrast, optogenetic inhibition of the striatum decreased the latency to SE onset in comparison to GFP control animals (Fig. 6B, p=0.0025, Mann-Whitney test).

**Figure 6.**
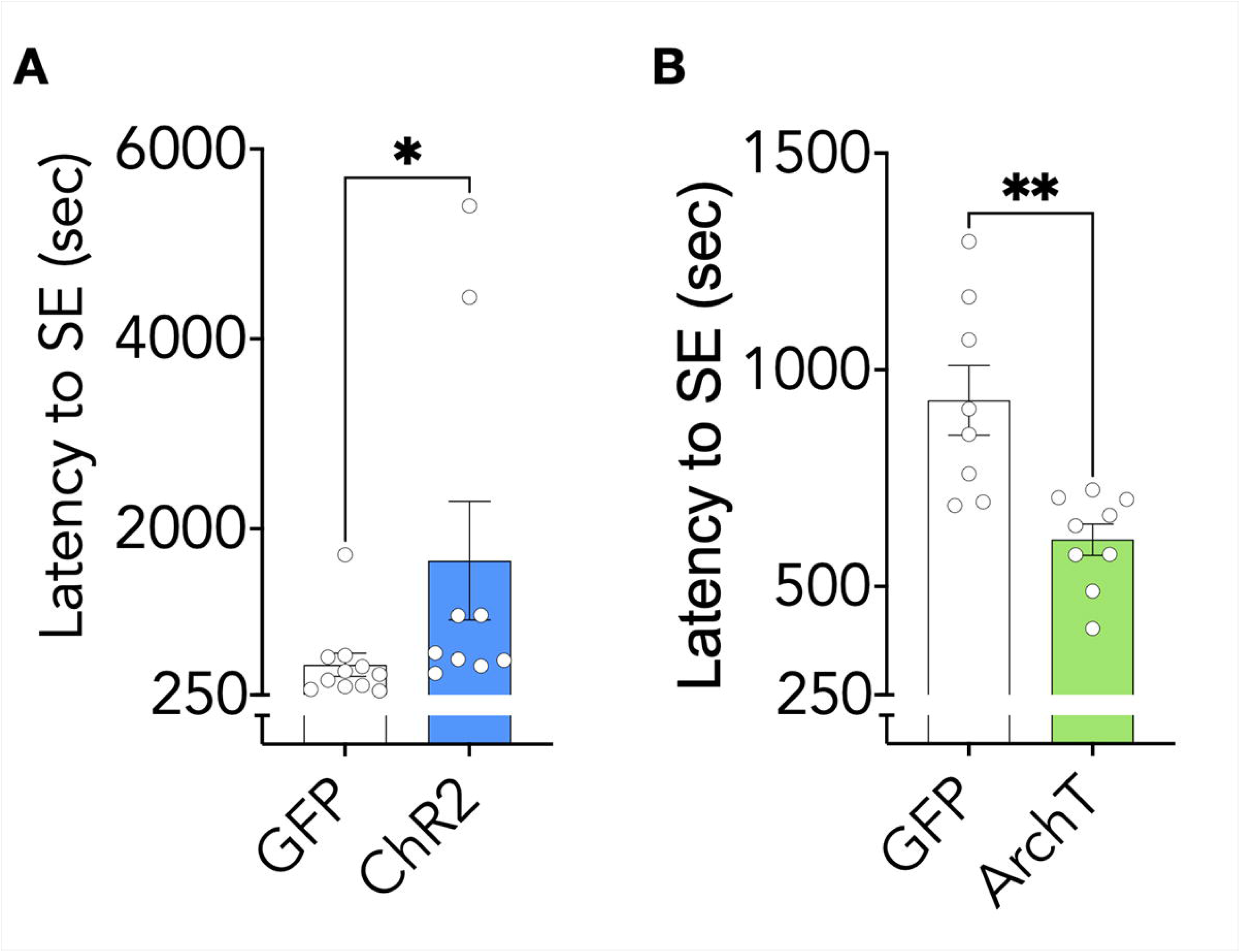
Optogenetic activation and inactivation of the dorsal striatum has bidirectional effects on pilocarpine-induced Status-Epilepticus (SE). (A-B) Optogenetic activation of the dorsal striatum increased the latency to SE onset in ChR2 animals (n=9) in comparison to GFP (no opsin) control rats (n=11) when both were stimulated with blue light at a frequency of 100 Hz. Optogenetic silencing of the dorsal striatum shortened the latency to SE onset in ArchT animals (n=9) compared to GFP control animals (n=8) when both groups were stimulated with green lights 100 Hz. Significance: p<0.05. *p<0.05, **p<0.01.

## Discussion

In this study, we report that the striatum displays a powerful bidirectional modulatory influence over seizures. We found that this effect is present across a range of seizure types, including absence seizures and two models of temporal lobe seizures, amygdala kindled seizures, and pilocarpine-induced SE. In the absence model we found that both open- and closed-loop neuromodulation strategies within the striatum were effective at modulating seizure activity. These data expand our knowledge of the extended basal ganglia system in the control of seizures and are consistent with the striatum’s role as the key upstream regulator of activity within the SNpr, a well-established anticonvulsant target (3, 4, 7, 22). Our data fill a clear gap: while striatal involvement in seizures has been known for 75 years, until now it was unclear if modulatory effects were present across seizure types due to a lack of comprehensive studies employing comparable interventions across multiple seizure models. Moreover, our data demonstrate that restricted activation of a small striatal portion – the dorsal striatum – is sufficient to modulate seizure activity. Relatively few studies have directly manipulated striatal activity in the context of seizures, however, in a series of studies in the 1980s, Turski and colleagues evaluated the effects of focal pharmacological manipulation in the striatum on limbic seizures. In these studies, they found that striatal activation via microinfusion of either bicuculline or NMDA protected against seizures in various models of limbic seizures (10, 11, 21). However, while the dorsal striatum of the rat is a relatively large structure, it lies close to crucial pathways that are known to be involved in limbic seizures. These include the structures within the limbic, frontal, and temporal cortices, as well as the nucleus accumbens (23–25). Drug diffusion in pharmacological microinjection studies is a challenge with respect to site-specificity. In the present study, we found that striatal optogenetic activation – which offers a much greater degree of site specificity – produced a marked protection against limbic seizures in response to electrical stimulation of the basolateral amygdala nucleus (BLA). Similarly, we also observed that striatal activation increased the latency to SE onset. Moreover, indicating a bidirectional impact of striatal activity on seizures, optogenetic inhibition of the striatum decreased the latency for pilocarpine-induced SE. The enhanced spatial specificity in the present study as compared to focal drug infusion in prior studies (which can be confounded by drug spread) provides strong evidence that these effects are truly striatally-mediated, and not due to off-target effects in other structures.

The effects of non-selective striatal activation and inhibition are somewhat paradoxical in the context of the standard model of basal ganglia function (22). D1 and D2 receptors are expressed on different populations of striatal output neurons, comprising the direct and indirect pathways, respectively. The D1 receptor is coupled to Gs signaling and expressed on GABAergic striatal neurons that project directly to the SNpr. Activation of D1 receptors thus is associated with increased activity in direct pathway neurons. By contrast, the D2 receptor is Gi-coupled and expressed on GABAergic neurons that project to the globus pallidus. GABAergic neurons in the globus pallidus in turn project to the subthalamic nucleus, which then sends excitatory glutamatergic projections to the SNpr. Activation of D2 receptors is thus associated with decreased activity in the indirect pathway, and a reduction in excitatory drive to the SNpr.

Based on extensive studies from many laboratories, inhibition of the SNpr is critical for basal ganglia-mediated anti-seizure effects (3, 4, 7, 26). How then, does non-selective striatal activation suppress seizures? In the standard model, one might predict a net neutral impact of striatal activation on SNpr activity. However, in a series of experiments performed using single unit recordings in the globus pallidus and SNpr, authors observed that ablation of striatal D2-expressing neurons (indirect pathway) had no impact on spontaneous firing rate of SNpr neurons (27). Consistent with this, while both GABA and glutamate iontophoresis modulates SNpr neurons in anesthetized animals, glutamate effects on SNpr neurons (mirroring activation of the indirect pathway) were limited in both magnitude and number of cells activated in awake rats. By contrast, in awake rats, GABA iontophoresis robustly inhibited firing. Consistent with this, electrical stimulation of the striatum robustly inhibits spontaneous firing activity in the SNpr, suggesting that non-selective striatal activation may have a more robust effect on the direct if compared to the indirect pathway (28). Together, these data suggest that GABA activity from the direct pathway seems to be the primary regulator of activity in SNpr neurons (29). Given that there are approximately 50 striatal direct pathway neurons for every SNpr neuron (30), and SNpr neurons receive dense synaptic input from direct pathway neurons, it is not surprising that direct pathway effects may predominate as a critical regulator of activity within the SNpr. It is also worth noting, however, that the standard model of basal ganglia function does not include pallido-nigral projections (31). Activation of pallidonigral GABAergic neurons, which collateralize to the STN, are also likely to trigger a pause in SNpr firing.

Only one study has indirectly examined the role of dopaminergic pathways in the ventral striatum (nucleus accumbens) in the context of absence seizures. Deransart and colleagues (32) found that combined injection of dopamine D1 and D2 receptor agonists in the accumbens reduced SWDs in the Genetic Absence Epilepsy Rat from Strasbourg (GAERS) model. By contrast, both D1 and D2 receptor antagonists worsened SWDs in the GAERS model. This is consistent with the general “accelerator” and “brake” model of the indirect and direct pathways. However, the effects of dopamine agonists and antagonists are modulators, rather than direct controllers of neuronal activity. We found that modulation of the striatal neurons elicited bidirectional control of SWDs: striatal activation suppressed SWDs and striatal inhibition worsened SWDs. Our data again suggest a predominant effect of the direct pathway on SWDs in the GBL model.

While we did not examine selective modulation of discrete striatal output pathways in the present study, non-selective activation, in some respects, is a more translational approach. Current deep brain stimulation methods are electrical and are unable to selectively activate divergent output pathways in a structure comprised of intermingled cell populations. From a network mechanism perspective, however, selective modulation of the direct and indirect pathway represents a clear next step. Likewise, while rodent models are valuable for our understanding of seizure modulatory networks, and while the gross organization of the basal ganglia is highly conserved across species, there are notable species differences in the organization of basal ganglia outputs. For example, while basal ganglia input regions, like the striatum and its projections, are fairly well-conserved (33, 34), the organization of output pathways from the substantia nigra differs. In the primate, SNpr neurons project to brainstem and thalamic targets with a low degree of collateralization (35, 36). By contrast in the rodent, nigral output neurons are highly collateralized (37). In some cases, the behavioral organization also differs across species (38, 39), underscoring the importance of future studies of the striatum as a DBS target in epilepsy, either in clinical populations, or in primate models.

In conclusion, our results demonstrate a role of the striatum as a modulator for seizures of various types including models of absence epilepsy and temporal lobe epilepsy, including both partial onset seizures and generalized onset seizures. The enhanced spatial specificity in the present study as compared to focal drug infusion in prior studies (which can be confounded by drug spread) provides strong evidence that these effects are truly striatally-mediated, and not due to off-target effects in other structures. Similarly, optogenetic neuromodulation allows for millisecond-scale on/off kinetics, allowing us to examine responsive neurostimulation, which produced robust effects on seizures. The ability to modulate seizures of various etiologies, seizures arising in distinct and discrete brain networks, and the ability to do so in a closed-loop manner point to the striatum as a potential translational target in patients with seizures of various types and in individuals with multifocal seizure onset as compared to conventional deep brain stimulation targets. Additionally, our present findings provide a strong rationale for future studies to separately examine the striatal direct and indirect pathways in the context of seizures.

## Materials and Methods

### Animals

92 adult male Sprague-Dawley rats were used. Animals were obtained from Charles River Laboratories (Wilmington, MA, USA) and were 3 months old at the beginning of the experiments. Two animals were housed per cage in a temperature and humidity-controlled room in the Division of Comparative Medicine (Georgetown University) with food (Lab Diet #5001) and water *ad libitum*. All experiments were performed during the light phase of the light/dark cycle (6:00 am. to 6:00 pm.). All procedures were approved by the Georgetown University Animal Care and Use Committee (Protocol 2016 1184).

### Surgery for virus injection and electrodes placement

For virus injection and electrodes placement, animals were anesthetized with ketamine (75 mg/kg; Zetamine; Vet Label) and dexmedetomidine (0.5 mg/kg; Dexdormitor; Orion Pharma Pfizer) and placed in a stereotaxic frame (Tujunga, CA).

For the GBL model and amygdala kindling models, animals received intra-striatal injection of AAV5-hsyn-ChR2 (H134R)-mCherry (opsin activation group), or AAV8-CAG-ArchT-GFP (opsin inactivation group), or AAV8-CAG-GFP (control, no opsin group). Both the synapsin and chicken B-actin promoters are expressed pan-neuronally (40).

All tests were performed using bilateral striatal neuromodulation. 1.5 μl of virus was bilaterally injected in the dorsal striatum at a rate of 0.2μl/min. The coordinates for the striatum were measured as follows on a flat skull plane: 1.7 mm anterior to the bregma, 2.5 mm lateral to the midline, and 4.5 mm ventral to the skull surface. After injection, fiber optics were bilaterally placed 0.2 mm above the site of injection.

For gamma-butyrolactone (GBL) and pilocarpine experiments, six cortical electrodes were implanted over the cortex, two over the frontal (2.5 mm anterior to the bregma and 2 mm lateral to the midline), two over the parietal (5 mm posterior to bregma and 2.5 mm lateral to the midline), and two (ground and reference) over the cerebellum (2.5 mm posterior to the lambda and 1 mm lateral to the midline). Frontal electrodes were used as reference to the cerebellum, the two parietal electrodes were referenced to each other. Electrodes wires were placed into a plastic pedestal (PlasticsOne, Roanoke, VA) and dental acrylic was used to hold and attach them to the skull.

For amygdala kindling experiments, a bipolar electrode was placed in the left hemisphere into the basolateral amygdala nucleus (BLA). The coordinates for the BLA were (again measured from a flat skull plane): 3.3 mm posterior to the bregma, 5.0 mm lateral to the midline, and 7.6 mm ventral to the skull surface. Four additional screws were placed over the parietal and frontal cortex to improve adherence of the headcap to the skull.

After surgery, animals were maintained in the Division of Comparative Medicine (DCM) for at least three weeks to allow sufficient time for opsin expression before experiments.

### Gamma-butyrolactone (GBL) model of generalized absence epilepsy

GBL (Sigma) was dissolved in saline 0.9% at a concentration of 100 mg/kg and intraperitoneally administered (1 ml/kg). The dose used was based on previous studies (41). At the beginning of each testing session, animals were tethered to a 3-channel recording system (Pinnacle Technologies, Lawrence, KS) via headstage containing cortical leads during surgery. Data were digitized and captured through a PowerLab interface and LabChart software (AD Instruments). After GBL injection, animals were monitored for spike-and-wave discharges (SWD) for 30 min in the open-loop (continuous light delivery) protocol of optogenetic neuromodulation and for 60 min in the closed-loop protocol (on-demand light-delivery). Each test session was performed with a minimum interval of 24 h.

The present study was designed based on within-subject comparisons, with testing order randomized and counterbalanced across sessions. This experimental design based on repeated measures and trials in the same animal minimizes the number of animals used in the study. Prior studies demonstrate that absence seizure duration is reduced when a second GBL administration was applied in the same day, but the same does not happen with 24 h between each administration (42).

### Amygdala kindling model of temporal lobe epilepsy

Stimulation in the amygdala kindling model was performed by an A-M Systems Isolated pulse stimulator (Model 2100) in constant current mode (A-M Systems, Sequim, WA). Electrical stimulations were delivered as a 1 s pulse of 60 Hz monophasic square wave pulse (1 msec pulse width). Electrographic responses were recorded by a kindling preamplifier with a solid-state relay to switch between record and stimulate modes (Pinnacle Technologies, Lawrence, KS). Signals were amplified (500 ×), digitized (1 KHz sampling rate), and filtered (band-pass 0.1 Hz – 30 Hz) (PowerLab, AD Instruments, Colorado Springs, CO) for offline analysis using LabChart Pro software (AD Instruments).

First, the afterdischarge threshold (ADT) was determined for each animal two weeks after surgery. ADT was defined as the minimum current intensity necessary to evoke spike activity that outlasted stimulation by at least 5 seconds. ADT determination began at a 20 μA current intensity, which was increased in steps of 20 μA every 2 min until the development of the ADT. After that, animals were daily stimulated at their specific pre-determined threshold. Seizures were analyzed according to the Racine score: 1: orofacial automatisms and ear myoclonus, 2: head myoclonus, 3: forelimb myoclonus, 4: forelimb myoclonus and elevation, 5: forelimb myoclonus, elevation, and fall (20).

Animals that displayed five consecutive Racine 5 were considered fully kindled, and fully kindled animals display highly stable responses to stimulation across weeks and months (43). Once animals were fully kindled a new ADT test was performed before the beginning of optogenetic stimulations to ensure animals were not tested using a supra-threshold current. For the experiments, animals were tested every 24h according to the experimental condition.

### Pilocarpine model of Status epilepticus (SE)

3 weeks after the last GBL session, animals were submitted to the pilocarpine-induced SE. Here, we did not use animals submitted to the amygdala kindling because kindling induces network changes that facilitate limbic seizures. Since we were unable to repeat SE in the same animals, we did not perform within-subject comparisons. All pilocarpine-induced SE were performed under continuous light delivery (open-loop).

Animals were pre-treated with scopolamine methylbromide (1 mg/kg/ml, ip.) to prevent effects associated with peripheral muscarinic activation. 30 min later, pilocarpine (380 mg/kg, 1ml/kg, ip.) was administered and EEG was recorded for 90 min. SE onset was defined as continuous (> 5 min) electrographic and behavioral seizures (Racine ≥ 3) and it was terminated with diazepam (10 mg/kg) 90 min after pilocarpine administration. Latency to SE onset was the dependent variable for this experiment.

### Optogenetic open-loop and closed-loop neuromodulation of the dorsal striatum

Fiber-optics were coupled to a fiber-coupled diode-pulsed solid-state laser (Doric Lenses; 450 nm for ChR2; 532 nm for ArchT) by a fiber-optic patch cord (fiber core 200 μm, NA 0.22). Laser power was calibrated before every test session with a power and energy meter to produce ∼10 mW out of the tip of the implanted fiber-optic. Stimulations were controlled with LabChart software (AD Instruments) and a PowerLab interface. Light was delivered at 5 or 100 Hz each with a 50% duty cycle. 5-Hz stimulation was delivered with a 100 ms pulse width and 100-Hz was delivered with 5 ms pulse width.

In the GBL model of absence epilepsy, animals were submitted to 4 open-loop test sessions (blue 5 Hz, blue 100 Hz, and two control sessions with no light delivery) and 2 closed-loop test sessions (blue 5 Hz and blue 100 Hz). In the open-loop protocol (continuous light delivery), optogenetic neurostimulation (5 or 100 Hz) was initiated immediately before GBL administration and lasted for the entire observation period. One control session was performed 24 hours before each optogenetic test condition, and no light was delivered during these sessions.

In the closed-loop protocol (on-demand light delivery), each animal was submitted to 2 test sessions (blue 5 Hz and blue 100 Hz). Seizures were detected in real time based on the EEG power and SWDs frequency. A macro, written in LabChart, performed an online fast Fourier transform (FFT) and calculated the power density (5–15 Hz). To minimize false positive SWD detections, seizure threshold was adjusted for each animal in every session based on EEG power and amplitude. The macro was written to deliver the light immediately after seizure detection. Seizures detected were randomized for two different conditions: light delivery or no light delivery (50% probability for each outcome). Lasers were adjusted to deliver 5 s of light for both light frequencies (5 and 100 Hz). To minimize overlapping seizure detection, a 10 s time-out was applied to the seizure detector after event detection.

In the amygdala kindling protocol we started optogenetic stimulation immediately prior the electrical stimulation and it lasted for the duration of the test session. Animals were submitted to two different test sessions with optogenetic stimulation (5 and 100 Hz) and two control sessions with no light delivery. Non-stimulated test sessions were performed 24h before optogenetic neuromodulation.

### Electroencephalography (EEG) recording and analysis

Prior to EEG recordings, animals were allowed to habituate in the experimental room for 20-30 min. Animals were placed in a transparent acrylic cylindric chamber and connected to a three-channel EEG headstage preamplifier connected to a commutator to the acquisition system.

EEG recordings were performed and acquired using a preamplifier-amplifier system from Pinnacle Technologies (Pinnacle 8206) and LabChart 8 (AD Instruments, Colorado Springs, CO). EEG was acquired at 10 kHz with a 1 Hz hardware highpass filter in the headstage preamplifier (10× head stage gain) and amplified (50×). Recordings were scored offline in LabChart by an observer blinded to the animal group and experimental test session. EEG recordings were filtered (bandpass 1–50 Hz) and SWDs were identified based on their amplitude (peak to peak exceeding 2x the background activity). The onset of the SWD was defined as the first peak in the discharge.

For optogenetic open-loop light delivery in the GBL-induced absence seizures, the number of SWDs, SWDs mean duration, and total seizure duration were measured. For the GBL closed-loop light delivery protocol, we measured the SWDs mean duration after light delivery. In the amygdala kindling protocol, the afterdischarge (AD) duration recorded from the BLA was calculated for each animal in every test session.

### Histological analysis

After the completion of the experiments, animals were overdosed with euthanasia solution (Euthasol, 390 mg pentobarbital sodium and 50 mg phenytoin sodium per ml) and perfused transcardially with phosphate buffered saline (PBS, 0.1 M, pH 7.4) followed by paraformaldehyde solution (PFA 4%, pH 7.4). Brains were removed, post fixed in PFA and cryoprotected prior to sectioning. Cryosections were stained with primary antibodies against GFP (rabbit polyclonal, 1:1000; ab290, abcam) or mCherry (1:1000; living colors DsRed, Takaha), with corresponding secondary antibodies and scanned using a Leica Mica confocal workstation.

### Statistical analysis

Statistical analyses were performed in GraphPad Prism (GraphPad Software, Inc, La Jolla, CA). Non-parametric data (e.g., seizure severity) were analyzed using Wilcoxon Matched Pairs test for paired data. Parametric data (e.g., SWDs duration) were tested for normality using the Shapiro-Wilk normality test and analyzed using paired t-tests. Unpaired data (e.g. latency to SE) were calculated using the Mann-Whitney test for non-parametric data and the Welch’s t-test to parametric data. Cumulative distribution of seizure duration in the closed-loop analysis was calculated by Kolmogorov–Smirnov test using the first 10 optogenetic stimulated and 10 non-stimulated seizures across rats within the same condition. The threshold for statistical significance was set at P < 0.05.

For absence epilepsy, a subset of recordings was scored by two observers, who reached a high rate of interobserver reliability for number of SWDs (R^2^ > 0.89), SWDs mean duration (R^2^ > 0.87), and time seized during test session (R^2^ > 0.91). For limbic seizures, behaviors were scored in real time and AD duration was calculated by two researchers (R^2^ > 0.87).

## Acknowledgments

This work was supported by R01NS097762 from the National Institutes of Health (NIH) / National Institute for Neurological Disorders and Stroke to PAF and F30NS110318 to SKH. We thank Dr. Carolina Campos-Rodriguez for helping with statistical analyses.

## References

1. J. Stoll, C. Ajmone-Marsan, H. H. Jasper, Electrophysiological studies of subcortical connections of anterior temporal region in cat. J. Neurophysiol. 14, 305–316 (1951).

2. G. B. Udvarhelyi, Dissemination of Acute Focal Seizures In the Monkey: I. From Cortical Foci. Arch. Neurol. 12, 333 (1965).

3. M. J. Iadarola, K. Gale, Substantia Nigra: Site of Anticonvulsant Activity Mediated by γ- Aminobutyric Acid. Science 218, 1237–1240 (1982).

4. D. S. Garant, K. Gale, Substantia nigra-mediated anticonvulsant actions: Role of nigral output pathways. Exp. Neurol. 97, 143–159 (1987).

5. P. Dean, K. Gale, Anticonvulsant action of GABA receptor blockade in the nigrotectal target region. Brain Res. 477, 391–395 (1989).

6. K. Morimoto, G. V. Goddard, The Substantia Nigra is an Important Site for the Containment of Seizure Generalization in the Kindling Model of Epilepsy. Epilepsia 28, 1–10 (1987).

7. E. Wicker, et al., Descending projections from the substantia nigra pars reticulata differentially control seizures. Proc. Natl. Acad. Sci. 116, 27084–27094 (2019).

8. C. Campos-Rodriguez, D. Palmer, P. A. Forcelli, Optogenetic stimulation of the superior colliculus suppresses genetic absence seizures. Brain awad166 (2023). 10.1093/brain/awad166.

9. C. Soper, E. Wicker, C. V. Kulick, P. N’Gouemo, P. A. Forcelli, Optogenetic activation of superior colliculus neurons suppresses seizures originating in diverse brain networks. Neurobiol. Dis. 87, 102–115 (2016).

10. L. Turski, et al., The basal ganglia, the deep prepyriform cortex, and seizure spread: bicuculline is anticonvulsant in the rat striatum. Proc. Natl. Acad. Sci. 86, 1694–1697 (1989).

11. E. A. Cavalheiro, L. Turski, Intrastriatal N-methyl-d-aspartate prevents amygdala kindled seizures in rats. Brain Res. 377, 173–176 (1986).

12. A. Brodovskaya, S. Shiono, J. Kapur, Activation of the basal ganglia and indirect pathway neurons during frontal lobe seizures. Brain 144, 2074–2091 (2021).

13. D. F. Aschauer, S. Kreuz, S. Rumpel, Analysis of Transduction Efficiency, Tropism and Axonal Transport of AAV Serotypes 1, 2, 5, 6, 8 and 9 in the Mouse Brain. PLoS ONE 8, e76310 (2013).

14. E. Krook-Magnuson, C. Armstrong, M. Oijala, I. Soltesz, On-demand optogenetic control of spontaneous seizures in temporal lobe epilepsy. Nat. Commun. 4, 1376 (2013).

15. B. J. Stieve, T. J. Richner, C. Krook-Magnuson, T. I. Netoff, E. Krook-Magnuson, Optimization of closed-loop electrical stimulation enables robust cerebellar-directed seizure control. Brain 146, 91–108 (2023).

16. K. Hristova, et al., Medial septal GABAergic neurons reduce seizure duration upon optogenetic closed-loop stimulation. Brain 144, 1576–1589 (2021).

17. Y. Takeuchi, et al., Closed-loop stimulation of the medial septum terminates epileptic seizures. Brain 144, 885–908 (2021).

18. E. Krook-Magnuson, G. G. Szabo, C. Armstrong, M. Oijala, I. Soltesz, Cerebellar Directed Optogenetic Intervention Inhibits Spontaneous Hippocampal Seizures in a Mouse Model of Temporal Lobe Epilepsy. eNeuro 1 (2014).

19. E. Wicker, P. A. Forcelli, Chemogenetic silencing of the midline and intralaminar thalamus blocks amygdala-kindled seizures. Exp. Neurol. 283, 404–412 (2016).

20. R. J. Racine, Modification of seizure activity by electrical stimulation: II. Motor seizure. Electroencephalogr. Clin. Neurophysiol. 32, 281–294 (1972).

21. L. Turski, et al., Dopamine-sensitive anticonvulsant site in the rat striatum. J. Neurosci. 8, 4027–4037 (1988).

22. C. Deransart, L. Vercueil, C. Marescaux, A. Depaulis, The role of basal ganglia in the control of generalized absence seizures. Epilepsy Res. 32, 213–223 (1998).

23. E. Valjent, G. Gangarossa, The Tail of the Striatum: From Anatomy to Connectivity and Function. Trends Neurosci. 44, 203–214 (2021).

24. S. Mraovitch, Y. Calando, Interactions between limbic, thalamo-striatal-cortical, and central autonomic pathways during epileptic seizure progression. J. Comp. Neurol. 411, 145–161 (1999).

25. S. J. Slaght, et al., Functional organization of the circuits connecting the cerebral cortex and the basal ganglia: implications for the role of the basal ganglia in epilepsy. Epileptic. Disord. 4, S9–S21 (2002).

26. S. Bröer, Not Part of the Temporal Lobe, but Still of Importance? Substantia Nigra and Subthalamic Nucleus in Epilepsy. Front. Syst. Neurosci. 14 (2020).

27. H. Sano, S. Chiken, T. Hikida, K. Kobayashi, A. Nambu, Signals through the Striatopallidal Indirect Pathway Stop Movements by Phasic Excitation in the Substantia Nigra. J. Neurosci. 33, 7583–7594 (2013).

28. M. Yoshida, N. Nakajima, K. Niijima, Effect of stimulation of the putamen on the substantia nigra in the cat. Brain Res. 217, 169–174 (1981).

29. F. Windels, E. A. Kiyatkin, GABA, Not Glutamate, Controls the Activity of Substantia Nigra Reticulata Neurons in Awake, Unrestrained Rats. J. Neurosci. 24, 6751–6754 (2004).

30. D. E. Oorschot, Total number of neurons in the neostriatal, pallidal, subthalamic, and substantia nigral nuclei of the rat basal ganglia: a stereological study using the cavalieri and optical disector methods. J. Comp. Neurol. 366, 580–599 (1996).

31. D. V. Simmons, M. H. Higgs, S. Lebby, C. J. Wilson, Indirect pathway control of firing rate and pattern in the substantia nigra pars reticulata. J. Neurophysiol. 123, 800–814 (2020).

32. C. Deransart, V. Riban, B.-T. Lê, C. Marescaux, A. Depaulis, Dopamine in the striatum modulates seizures in a genetic model of absence epilepsy in the rat. Neuroscience 100, 335–344 (2000).

33. C. D. Hardman, et al., Comparison of the basal ganglia in rats, marmosets, macaques, baboons, and humans: Volume and neuronal number for the output, internal relay, and striatal modulating nuclei. J. Comp. Neurol. 445, 238–255 (2002).

34. W. J. a. J. Smeets, O. Marín, A. González, Evolution of the basal ganglia: new perspectives through a comparative approach. J. Anat. 196, 501–517 (2000).

35. R. M. Beckstead, A. Frankfurter, The distribution and some morphological features of substantia nigra neurons that project to the thalamus, superior colliculus and pedunculopontine nucleus in the monkey. Neuroscience 7, 2377–2388 (1982).

36. A. Parent, A. Mackey, Y. Smith, R. Boucher, The output organization of the substantia nigra in primate as revealed by a retrograde double labeling method. Brain Res. Bull. 10, 529–537 (1983).

37. C. Cebrián, A. Parent, L. Prensa, Patterns of axonal branching of neurons of the substantia nigra pars reticulata and pars lateralis in the rat. J. Comp. Neurol. 492, 349–369 (2005).

38. B. L. Aguilar, P. A. Forcelli, L. Malkova, Inhibition of the substantia nigra pars reticulata produces divergent effects on sensorimotor gating in rats and monkeys. Sci. Rep. 8, 9369 (2018).

39. D. Dybdal, et al., Topography of dyskinesias and torticollis evoked by inhibition of substantia nigra pars reticulata. Mov. Disord. Off. J. Mov. Disord. Soc. 28, 460–468 (2013).

40. M. Zhao, R. Alleva, H. Ma, A. G. S. Daniel, T. H. Schwartz, Optogenetic tools for modulating and probing the epileptic network. Epilepsy Res. 116, 15–26 (2015).

41. V. R. Santos, I. Kobayashi, R. Hammack, G. Danko, P. A. Forcelli, Impact of strain, sex, and estrous cycle on gamma butyrolactone-evoked absence seizures in rats. Epilepsy Res. 147, 62–70 (2018).

42. P. K. Banerjee, et al., Alterations in GABAAReceptor α1 and α4 Subunit mRNA Levels in Thalamic Relay Nuclei Following Absence-like Seizures in Rats. Exp. Neurol. 154, 213–223 (1998).

43. Z. Dennison, G. Campbell Teskey, D. P. Cain, Persistence of kindling: Effect of partial kindling, retention interval, kindling site, and stimulation parameters. Epilepsy Res. 21, 171– 182 (1995).

44. G. Paxinos, C. Watson, The Rat Brain in Stereotaxic Coordinates: Hard Cover Edition (Elsevier, 2006).

