## Supplemental Figures for "Optogenetic stimulation of dorsal striatum bidirectionally controls seizures"

### Frontal cortex - Left

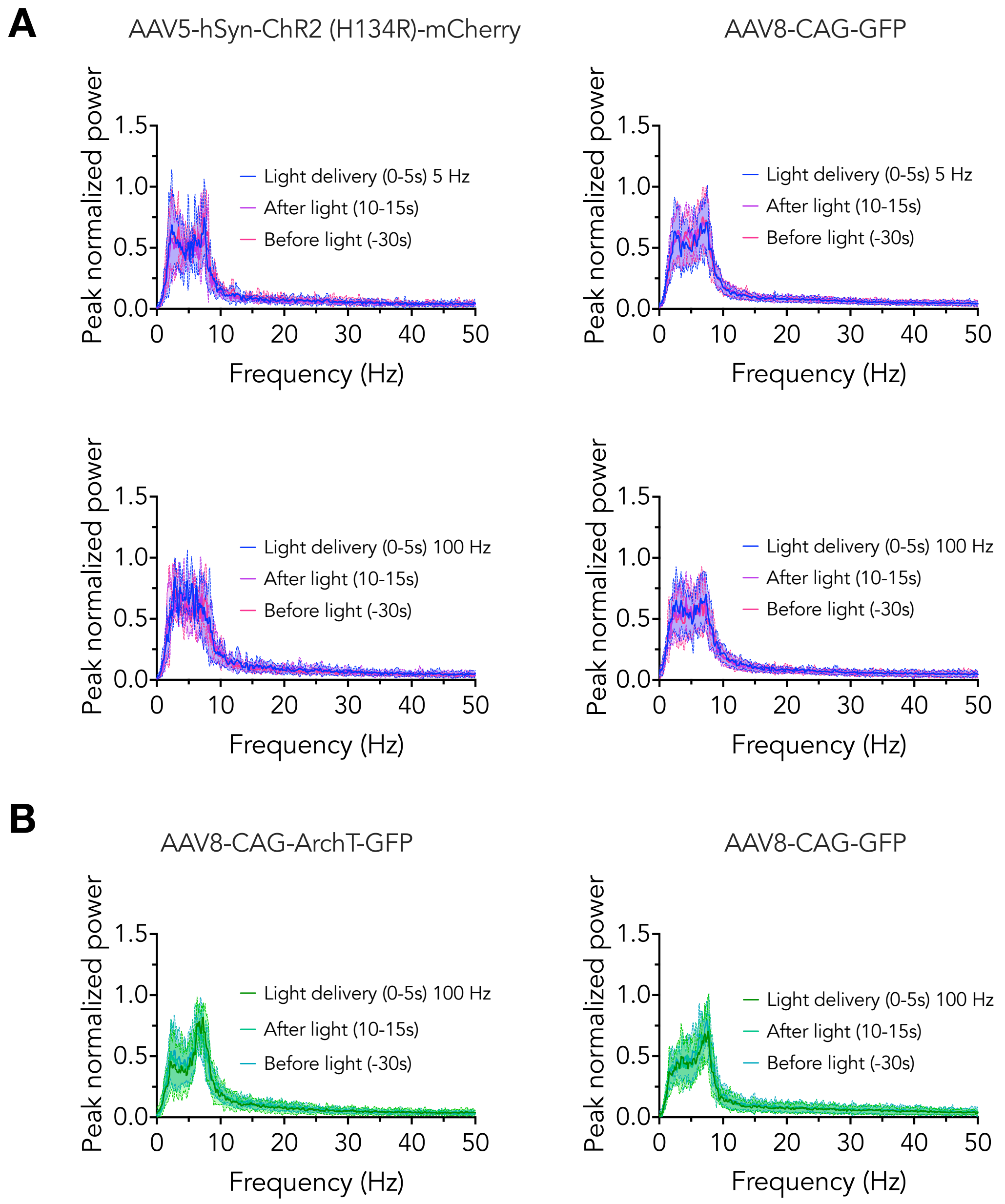


**Figure S1.** Bilateral optogenetic stimulation of the dorsal striatum did not change power spectra activity, measured by peak normalized power, on left frontal cortex. (A) 5 seconds of blue light delivery (5 and 100 Hz; 0-5s) did not change EEG power spectra compared to both 30 seconds before light delivery and 5 seconds after light delivery. (B) 5 seconds of green light delivery (100 Hz; 0-5s) did not change EEG power spectra compared to both 30 seconds before light delivery and 5 seconds after light delivery. AAV5-hSyn-ChR2 (h134R)-mCherry – striatal activation group. AAV8-CAG-ArchT-GFP – striatal inactivation group. AAV8-CAG-GFP – no opsin control group.

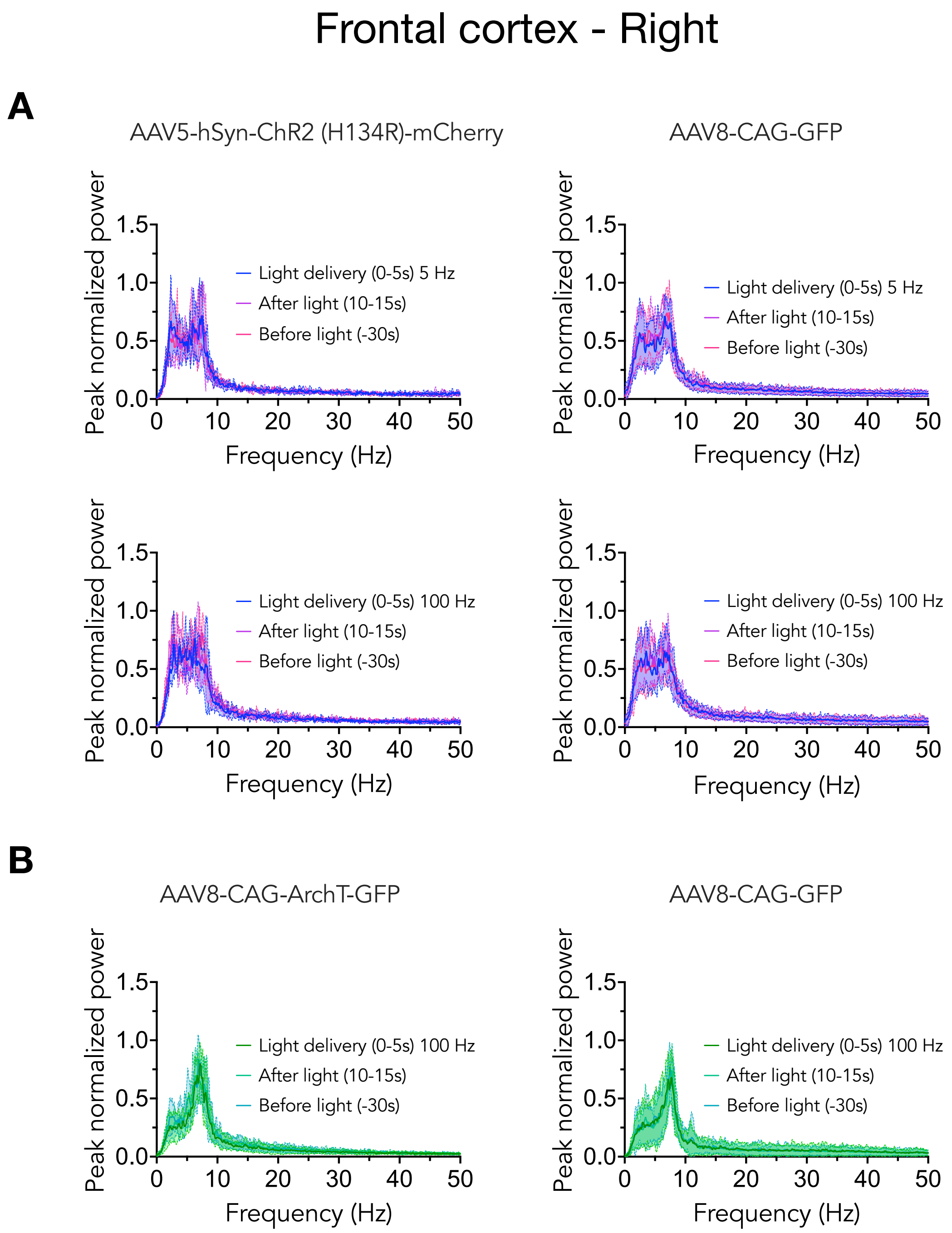


**Figure S2.** Bilateral optogenetic stimulation of the dorsal striatum did not change power spectra activity, measured by peak normalized power, on right frontal cortex. (A) 5 seconds of blue light delivery (5 and 100 Hz; 0-5s) did not change EEG power spectra compared to both 30 seconds before light delivery and 5 seconds after light delivery. (B) 5 seconds of green light delivery (100 Hz; 0-5s) did not change EEG power spectra compared to both 30 seconds before light delivery and 5 seconds after light delivery. AAV5-hSyn-ChR2 (h134R)-mCherry – striatal activation group. AAV8-CAG-ArchT-GFP – striatal inactivation group. AAV8-CAG-GFP – no opsin control group.

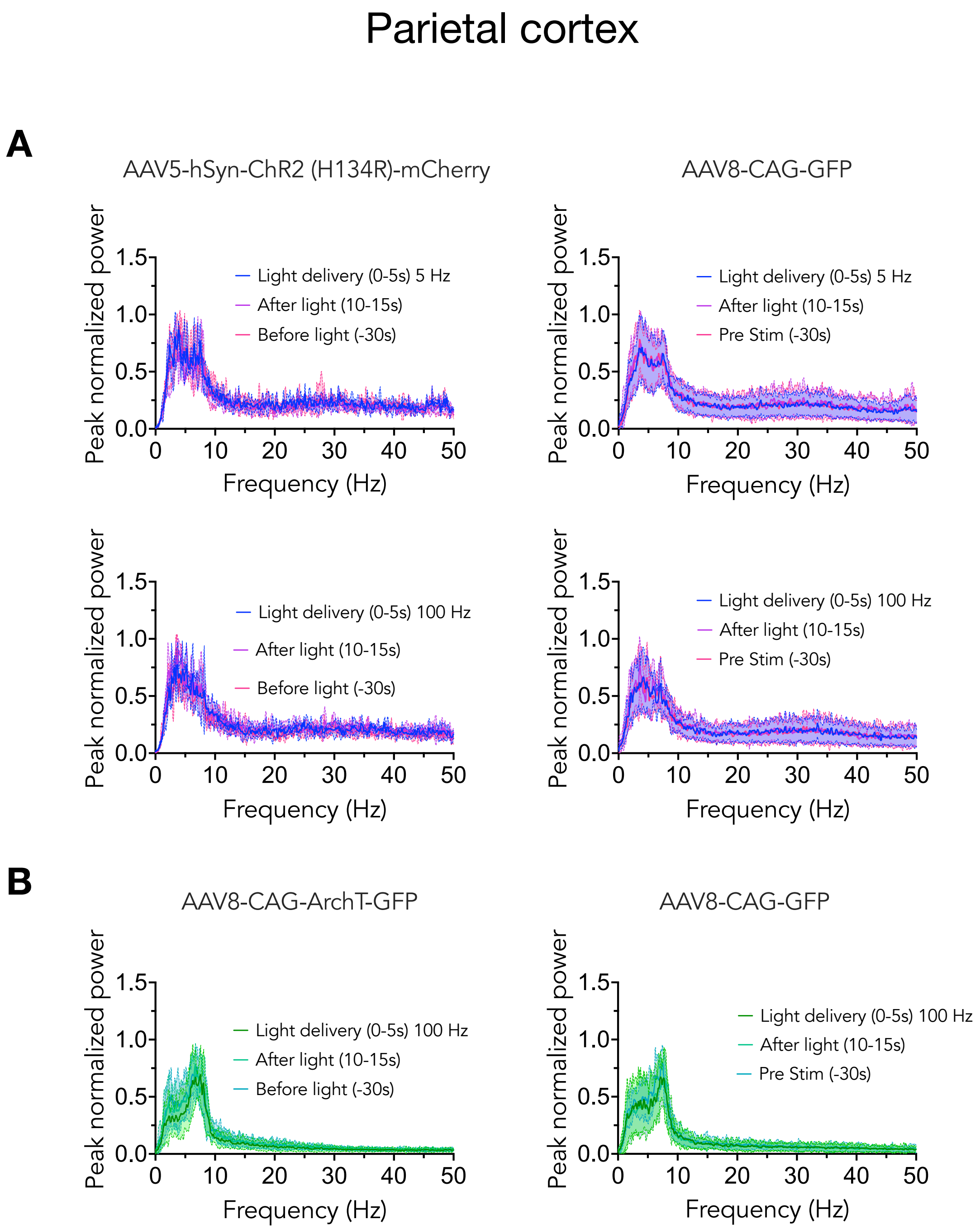


**Figure S3.** Bilateral optogenetic stimulation of the dorsal striatum did not change power spectra activity, measured by peak normalized power, on parietal cortex. (A) 5 seconds of blue light delivery (5 and 100 Hz; 0-5s) did not change EEG power spectra compared to both 30 seconds before light delivery and 5 seconds after light delivery. (B) 5 seconds of green light delivery (100 Hz; 0-5s) did not change EEG power spectra compared to both 30 seconds before light delivery and 5 seconds after light delivery. AAV5-hSyn-ChR2 (h134R)-mCherry – striatal activation group. AAV8-CAG-ArchT-GFP – striatal inactivation group. AAV8-CAG-GFP – no opsin control group.

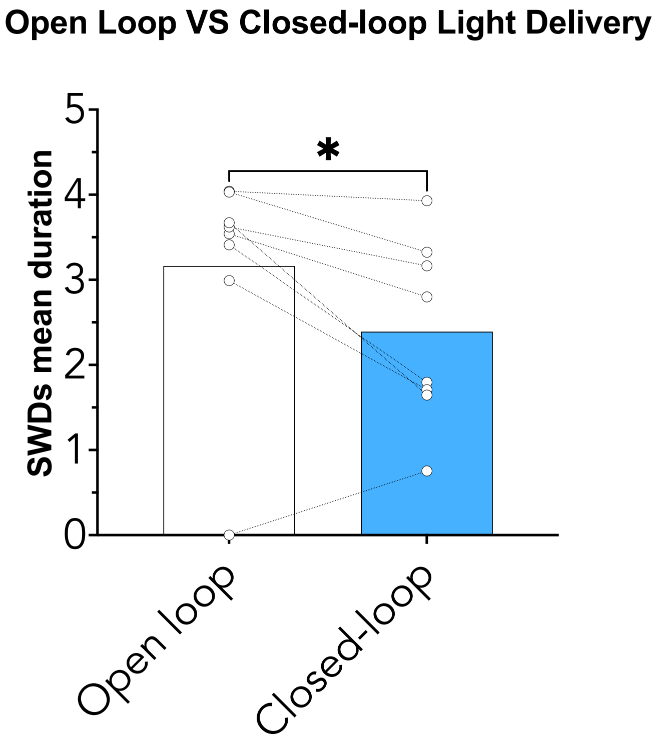


**Figure S4.** Dorsal striatal closed-loop light (blue 100 Hz) delivery suppress SWDs faster than in the open-loop protocol (blue 100 Hz) in ChR2-expressing animals. N=8. Significance: $p<0.05$.

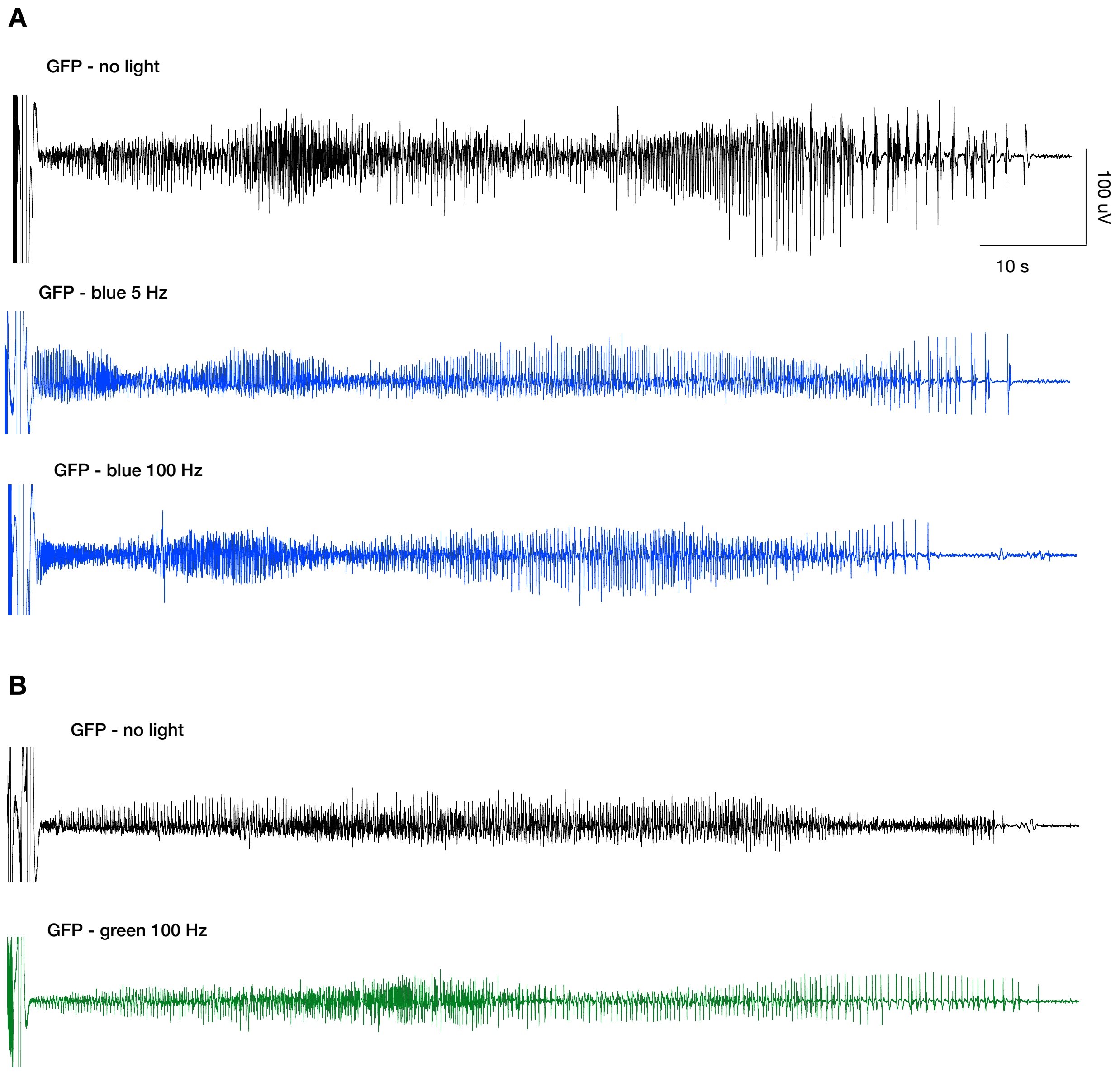


**Figure S5.** Representative EEG trace recorded from the basolateral amygdala nucleus (BLA) of a GFP control animal (AAV8-CAG-GFP, opsin negative) during no light (black traces), blue light delivery (5 and 100 Hz) and green light delivery (100 Hz). Sessions were performed after the animal were fully kindled. No significant difference was detected regarding afterdischarge (AD) duration across the sessions.

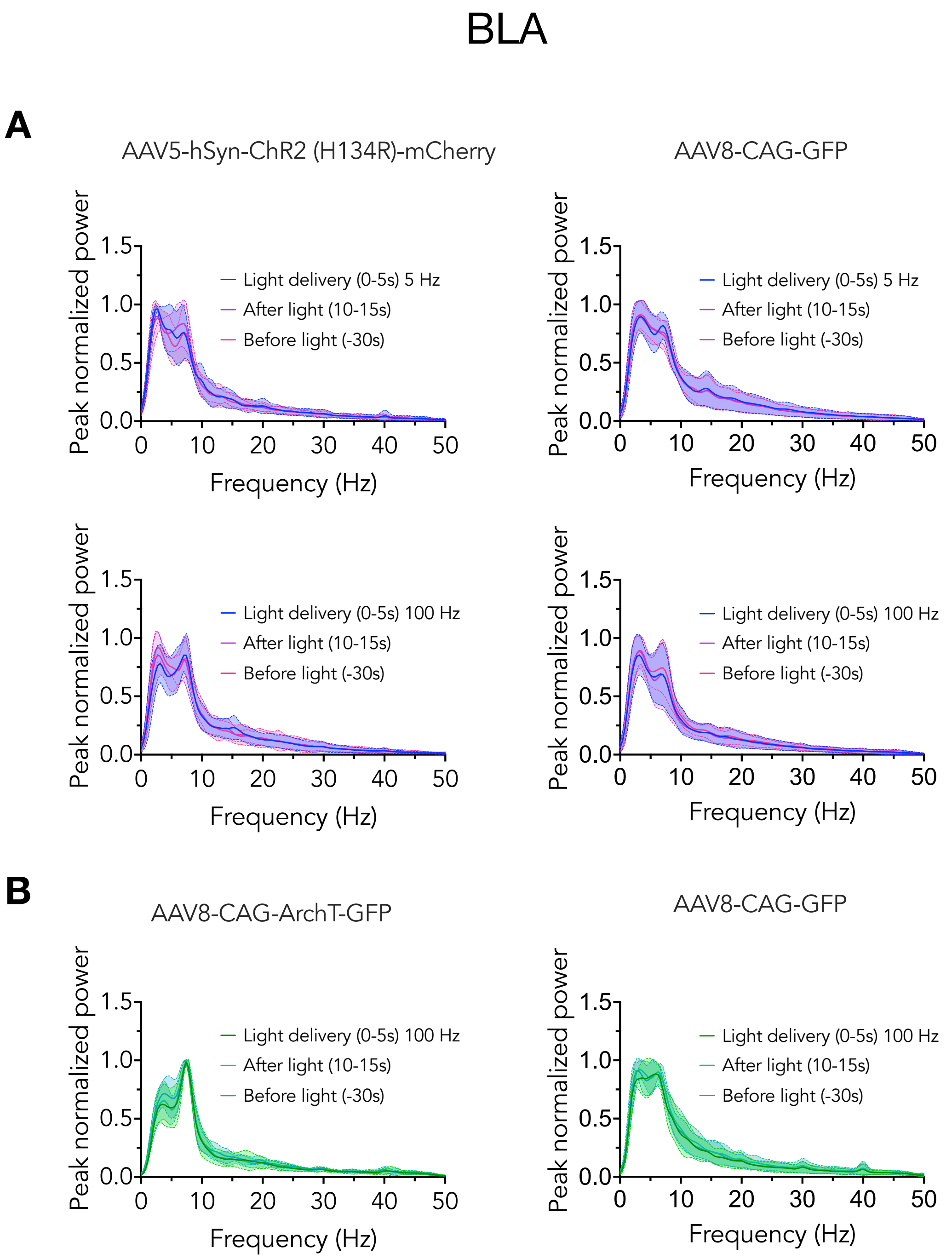


**Figure S6.** Bilateral optogenetic stimulation of the dorsal striatum did not change power spectra activity, measured by peak normalized power, on basolateral amygdala nucleus (BLA). (A) 5 seconds of blue light delivery (5 and 100 Hz; 0-5s) did not change EEG power spectra compared to both 30 seconds before light delivery and 5 seconds after light delivery. (B) 5 seconds of green light delivery (100 Hz; 0-5s) did not change EEG power spectra compared to both 30 seconds before light delivery and 5 seconds after light delivery. AAV5-hSyn-ChR2 (h134R)-mCherry –

striatal activation group. AAV8-CAG-ArchT-GFP – striatal inactivation group. AAV8-CAG-GFP – no opsin control group.
